# Using hospital network-based surveillance for antimicrobial resistance as a more robust alternative to self-reporting

**DOI:** 10.1101/535252

**Authors:** Tjibbe Donker, Timo Smieszek, Katherine L. Henderson, Timothy M. Walker, Russell Hope, Alan P. Johnson, Neil Woodford, Derrick W. Crook, Tim E.A. Peto, A. Sarah Walker, Julie V. Robotham

**Author notes:** Corresponding author: Tjibbe Donker, Nuffield Department of Medicine, University of Oxford, John Radcliffe Hospital, Oxford, OX3 9DU, UK. These authors contributed equally (Tim E.A. Peto, A. Sarah Walker, Julie V. Robotham).

## Abstract

Hospital performance is often measured using self-reported statistics, such as the incidence of hospital-transmitted micro-organisms or those exhibiting antimicrobial resistance (AMR), encouraging hospitals with high levels to improve their performance. However, hospitals that increase screening efforts will appear to have a higher incidence and perform poorly, undermining comparison between hospitals and disincentivising testing, thus hampering infection control. We propose a surveillance system in which hospitals test patients previously discharged from other hospitals and report observed cases. Using NHS Hospital Episode Statistics data, we analysed patient movements across England and assessed the number of hospitals required to participate in such a reporting scheme to deliver robust estimates of incidence. With over 1.2 million admissions to English hospitals previously discharged from other hospitals annually, even when only a fraction of hospitals (41/155) participate (each screening at least 1000 of these admissions), the proposed surveillance system can estimate incidence across all hospitals. By reporting on other hospitals, the reporting of incidence is separated from the task of improving own performance. Therefore the incentives for increasing performance can be aligned to increase (rather than decrease) screening efforts, thus delivering both more comparable figures on the AMR problems across hospitals and improving infection control efforts.

## Introduction

Many healthcare systems worldwide mandate the reporting of key hospital statistics to measure performance[1]. Such self-reported assessments are intended to provide a clear, comparable overview of each hospital’s status, by ranking them based on their reported statistics. Poorly performing hospitals can then be encouraged to improve using incentives ranging from financial penalties[2,3] to reputational damage through ‘naming and shaming’. The mandatory reporting of antimicrobial resistance (AMR) and other hospital-transmitted organisms are examples of commonly used self-reporting systems[4].

Surveillance systems for AMR are attractive to policy-makers, as they can be used to increase patient safety by identifying where extra infection prevention and control (IPC) efforts need to be coordinated, as well as providing insight into the spread and epidemiology of AMR. Changes in incidence after introducing such systems, like the dramatic decline in methicillin-resistant Staphylococcus aureus (MRSA) bacteraemia after the initiation of the mandatory surveillance scheme in the United Kingdom[5], have led some to conclude that such self-reporting surveillance systems help reduce rates.

However, true incidence of AMR is often hard to measure, because large numbers of affected patients may be asymptomatically colonised[6,7] and thus only found when actively screened. Hospitals targeting screening strategies to identify more cases may thus worsen their ranking by increasing their reported incidence. Systems of assessing hospitals based on self-reported carriage rates may thus unintentionally punish hospitals with stringent testing, screening, and reporting regimes, because of their seemingly poor performance. Both IPC efforts and hospital performance monitoring may therefore be hindered by the conflicting incentives: to improve IPC efforts, a hospital needs to identify as many cases as possible, while it needs to find as few as possible to improve its performance ranking.

We explore how to align incentives for hospitals, by separating the task of reporting incidence of a predominantly carried micro-organism that is acquired in hospital from the task of lowering its incidence. To do this, we propose a novel surveillance system based on the hospital network formed by shared patients, namely testing patients that were previously admitted to another hospital to provide an approximation of the incidence of AMR in that hospital. We show the potential of this network-based surveillance system to provide incidence estimates, and explore its operational limits, in particular the number of participating hospitals needed to reliably estimate incidences for all hospitals. We argue that such a system can provide a more robust surveillance system for AMR than self-reporting.

## Methods

### Network-based surveillance system

In the proposed surveillance system (Figure 1), each hospital reports the number of patients previously admitted to and discharged from other hospitals in a predefined time-frame (e.g. the previous 12 months) and found to be colonised when screened on admission to this index hospital (denoted imported cases). The reported numbers of imported cases are then pooled to give the total number of found cases exported from all the hospitals across the network. For simplicity of reporting, any untested patients are assumed to not be colonised (providing an incentive to test admissions previously discharged from elsewhere). The number of imported cases are then divided by the total number of patients previously discharged from that hospital and admitted to one of the reporting hospital (which can be obtained from central statistics) to give an estimate of incidence. Alternatively, without loss of generality, numbers tested and testing positive could be reported and summed to give an estimate of incidence.

**Figure 1.**
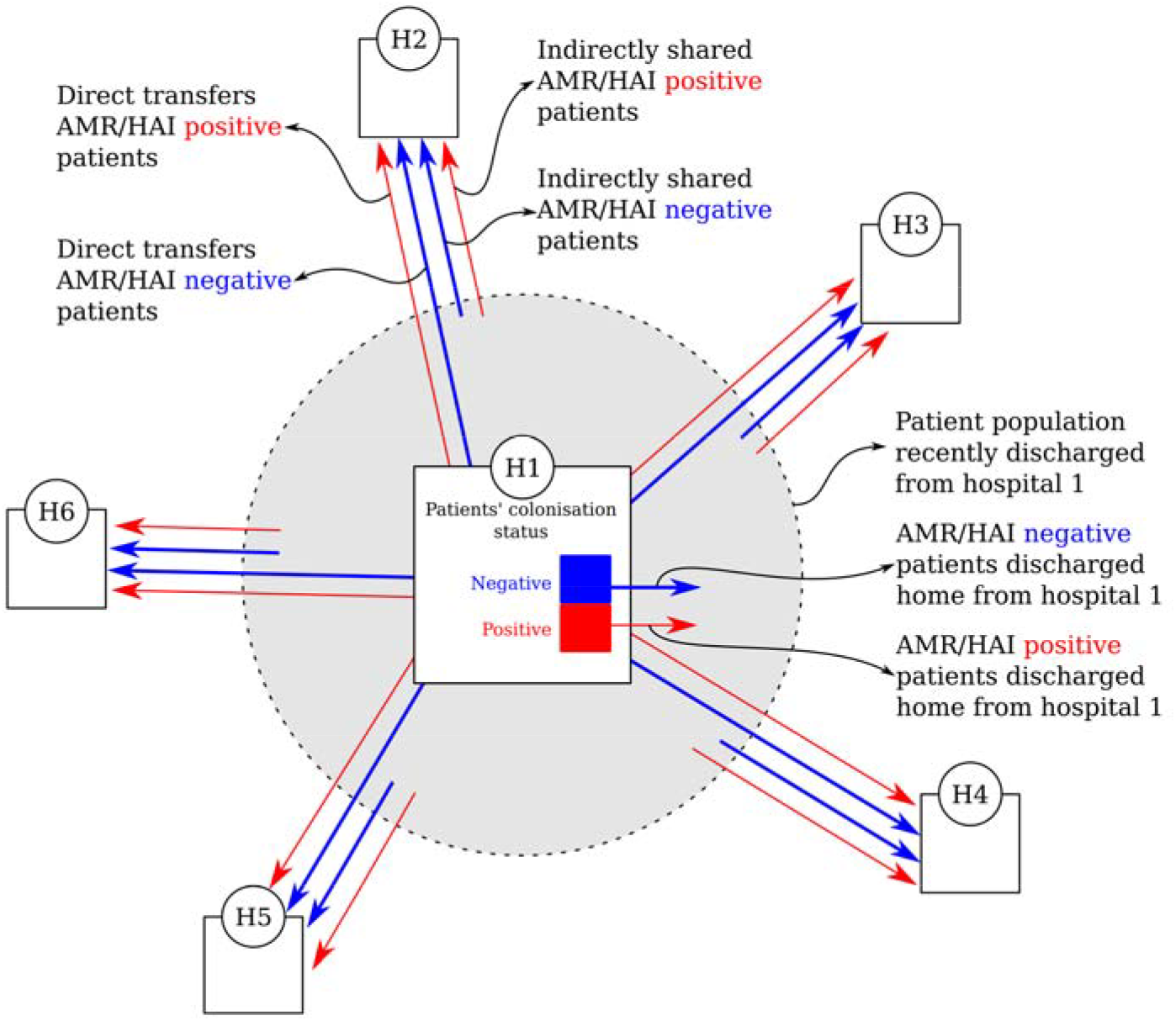
Schematic representation of the proposed surveillance system. A proportion of the patients discharged from hospital 1 will be directly transferred or indirectly readmitted to hospitals 2-6. These shared patients may carry AMR acquired in hospital 1. By reporting these colonised patients, as well as the total number of shared patients, hospitals 2-6 can estimate an AMR incidence for hospital 1 without hospital 1 reporting.

To demonstrate, we use data on patient admissions from the National Health Service (NHS) Hospital Episode Statistics (HES) to determine the number of patients that were admitted to different English hospital Trusts (denoted ‘hospitals’) post discharge. We sorted all admissions per patient by admission date; and for all admissions during 2014-’15 determined whether the previous discharge happened 1 year before the admission date and whether the previous admission was to a different hospital to the current one (i.e. the patient was shared between hospitals). Sensitivity analyses considered 6, 3, 1 month and one week.

Each hospital had two sets of admissions associated with it: 1) all admissions (the general patient population), and 2) a subset the admissions of patients previously discharged from another hospital, now admitted to this hospital (the received patients). The received patient population comes from a number (potentially all) of the other hospitals. We therefore denote the number of patients discharged from hospital *i* and subsequently admitted to hospital *j* as m_ij_, where s_i_=Σj m_ij_ is the total shared population size from hospital *i*. Under the proposed surveillance scheme, these received patients should be screened as they are admitted to hospital *j* to gather information about the incidence of hospital-associated pathogens in hospital *i*.

### Coverage

The system consists of the “reporting set”, namely hospitals reporting the number of AMR cases among their received patients, and the “covered set”, namely hospitals whose discharged patients are screened as they arrive in other hospitals. We consider a hospital to be part of the covered set once a fixed number of its discharged patients per year (the reporting threshold) are received by the hospitals within the reporting set. Thus the reporting set does not necessarily need to include all hospitals for the covered set to include all hospitals.

Any hospital sharing fewer patients than this reporting threshold with all other hospitals combined cannot, by definition, be reported on by such a scheme. Thus the minimum number of patients shared by hospitals is the highest reporting threshold that can be used (n=1216). Taking 1000 shared patients as the reporting threshold, we determined the total number of hospitals that need to be included in the surveillance scheme to be able to report on all hospitals in three ways; first by random assignment, second by adding hospitals based on the number of received patients, and third by adding hospitals using a greedy algorithm.

### Assignment of hospitals

For the first selection procedure, we randomly added hospitals to the reporting set, one at a time, calculating the number of hospitals in the covered set after each addition. Hospitals were added to the reporting set until all hospitals were included in the covered set, repeating this algorithm 100 times. For the second procedure (receipt-based), we sorted hospitals based on the total number of patients they received from other hospitals, and added them to the reporting set, starting with the hospital that received most patients and iteratively adding the other hospitals to maximise the number of received patients added at each step.

The greedy algorithm iteratively added the hospital to the reporting set that would add the most hospitals to the covered set. Per step, we calculated for each reporting hospital how many other hospitals it would add information on (i.e. by how many hospitals the covered set would increase if this hospital was added to the reporting set). If the number of covered hospitals did not increase by adding any hospital, the hospital that resulted in the largest increase in number of received patients from hospitals not yet included in the covered set was added. The same procedure was used if two hospitals resulted in the same increase to the covered set.

### Reciprocal reporting (snow-ball effect)

We further tested the effect of assuming that covered hospitals will automatically start reporting once they are themselves reported on, based on the game-theoretical considerations that hospitals will try to ‘win’ the ranking of reported incidences (supplementary text). After adding a hospital following the greedy algorithm, we checked if all covered hospitals were present in the reporting set and added them if they were not. Because the increase in reporting could increase the number of covered hospitals, this step was repeated until no hospitals were added to the reporting and covered sets. After this, the next hospital was added to the reporting set using the greedy algorithm again.

## Results

### Network-based surveillance

To test the feasibility of having hospitals report the number of patients previously admitted to other hospitals that are AMR (or other equivalent carried micro-organism) positive on admission, rather than self-reporting their own patients colonised on or during admission, we reconstructed the English hospital network (Figure 2A), based on the NHS Hospital Episode Statistics for England. The network consisted of 155 hospital organisations (so-called Trusts, denoted ‘hospitals’ for generalisability) during the financial year 2014-15, admitting 8,681,397 patients for a total of 15,708,764 admissions. A total of 1,208,999 admissions were preceded within a year by a discharge from a different hospital, mainly concentrated within a small number of strong connections between hospitals (Figure 2B). The median time between the previous discharge and admission was 28 days (IQR 6-104), the mean number of overnight stays was 2.1 (IQR 0-2, median 0) for all patient admissions (Figure 2C), while shared patients stayed 4.6 nights (IQR 0-4, median 1).

**Figure 2.**
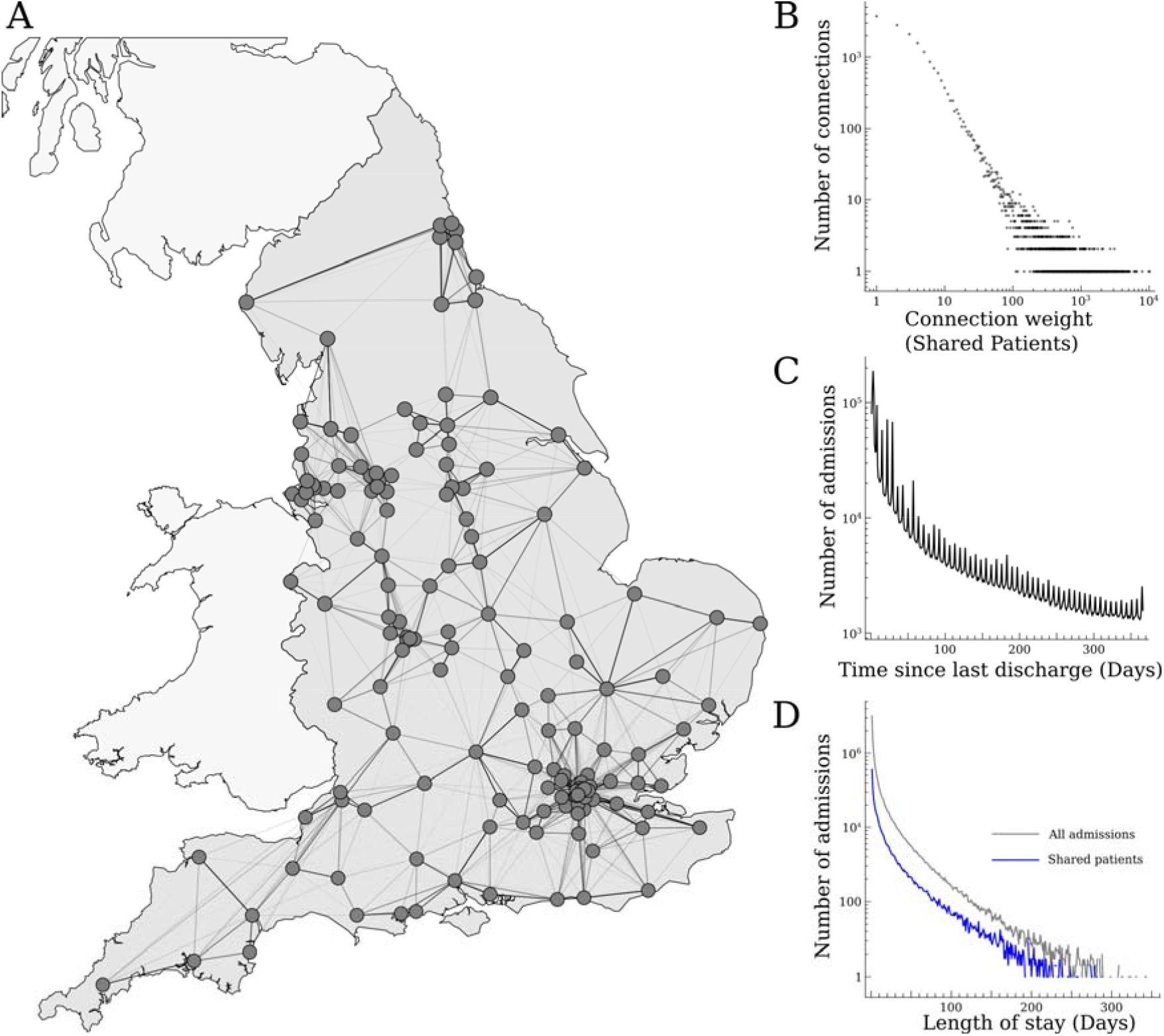
The English hospital network. A) The location of the included hospitals (dots), showing the connections and connection weights based on patients shared between them (admitted to one hospital having previously been discharged from another) (lines, darkness indicating the number of shared patients). B) The distribution of connection weights between all hospitals. C) The distribution of time between admissions, measured as days since previous discharge. D) The distribution of lengths of stay, for all admissions (grey) and shared patients (blue).

The number of shared patients (patients who were first admitted to a certain hospital, and subsequently admitted to any of the others) was highest for a tertiary care hospital in the North-East (23,260 received by others), and lowest for a cancer centre in the North-West (1,216 received by others). Based on 1,216 as the upper limit of patients that can be received from the least connected hospital, we set our reporting threshold at 1000. If the maximum time between discharge and subsequent admission was reduced from a year to a week, the number of subsequent admissions was reduced by about 78% (Figure 3A), with a total of 264,920 subsequent admissions, of which 5,314 were received from the most-connected (a London teaching hospital) and 232 from the least-connected (an orthopaedic hospital). Specialist hospitals shared the fewest patients, and higher thresholds up to 2,989 can be used to include the remaining 146 hospitals when these nine specialists are excluded.

**Figure 3.**
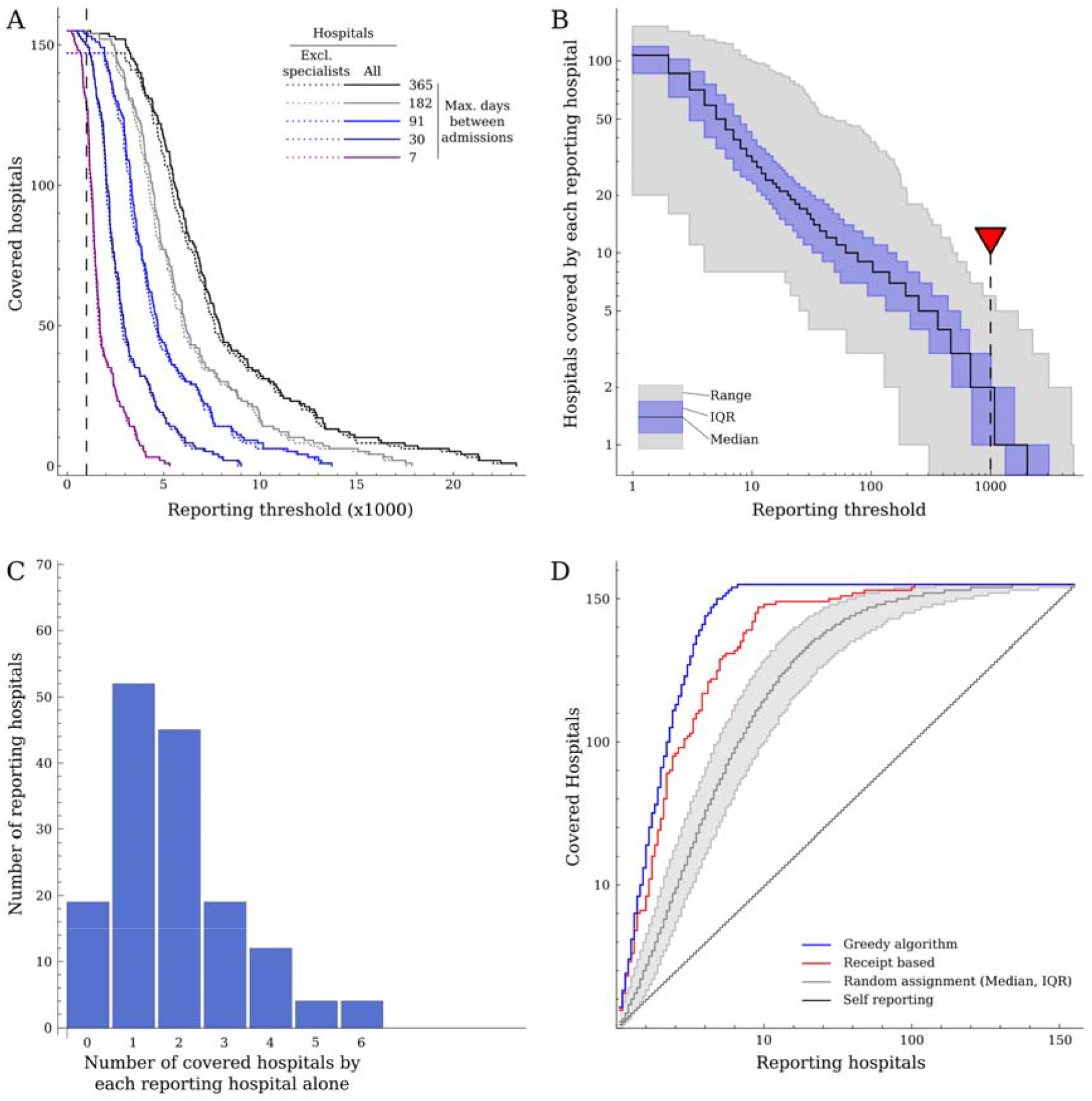
A) The number of patients discharged from each hospital and subsequently admitted elsewhere for different maximum periods between last discharge and next admission. If previous discharges within a year are included, all hospitals discharge over 1000 patients who are subsequently admitted elsewhere within a year. B) The number of hospitals that are covered by each reporting hospital individually, as a function of the threshold number of received patients. C) The number of hospitals that are covered by each reporting hospital individually, for a threshold of 1000 received patients (shown by red triangle in B). D) The number of hospitals covered as a function of the number of reporting hospitals using self-reporting (black line) as well as the proposed surveillance scheme with the reporting set determined by random assignment (grey), receipt-based assignment (blue) and the greedy algorithm (blue).

A key feature of this system is that hospitals can be included in the covered set even if none of the individual reporting hospitals receive over the threshold of 1000 patients, as long as all hospitals combined receive over this threshold. In fact, a median 134 hospitals (of total 155) were required in a randomly chosen reporting set to provide enough data to include information about all hospitals in the covered set. Strikingly, a median of only 30 hospitals needed to be included in a randomly chosen reporting set to survey incidence in half (n=78) of the hospitals. Numerous hospitals received enough patients to be able to individually report on several others (Figure 3B). Four hospitals each reported on six other hospitals at the 1000-patient threshold (Figure 3C). The number of hospitals in the covered set (achieving the threshold of >1000 received patients) was always higher than the number of reporting hospitals (Figure 3D). In contrast, and by definition, any self-reporting scheme reports only on exactly the numbers of hospitals included in the scheme.

By selecting hospitals into the reporting set based on the number of patients they received from other hospitals (labelled “receipt-based” in Figure 3D), the reach of the covered set could be substantially improved, with incidence estimated from >1000 patients in half the hospitals after including just 16 hospitals in the reporting set. However, to estimate incidence in all hospitals, this selection procedure still needed to include 101 hospitals in the reporting set.

A “greedy” algorithm significantly outperformed both the random and receipt-based additions to the reporting set, increasing the covered set faster and providing the largest number of covered hospitals (with incidence estimated from >1000 patients) for any number of reporting hospitals. The difference between the greedy algorithm and the receipt-based selection was largest for the last 50 covered hospitals. Incidence could be estimated from >1000 patients in all hospitals after adding only 41 hospitals to the reporting set using the greedy algorithm (Figure 3D & 4A), while only 13 reporting hospitals were needed to survey 50% of all hospitals.

In the so-called “snowball” scenario, where hospitals start reporting if they are reported on, the number of reporting hospitals quickly expands. After the first hospital starts reporting its received cases, its neighbours will join, followed by their neighbours, each time increasing the number of received cases that are reported and the likelihood of other hospitals adding themselves to the covered set (Figure 4B). For most randomly selected starting hospitals, this resulted in all hospitals eventually being included in the reporting set. Only if the first hospital was small enough to not receive >1000 patients from any particular hospital did the first step not result in the addition of more hospitals to the reporting set (occurring with probability 19/155=0.12). For a group of nine hospitals in the North, the snowball-addition stopped when the whole group was added, as the nine hospitals combined did not receive >1000 patients from any other hospitals.

**Figure 4.**
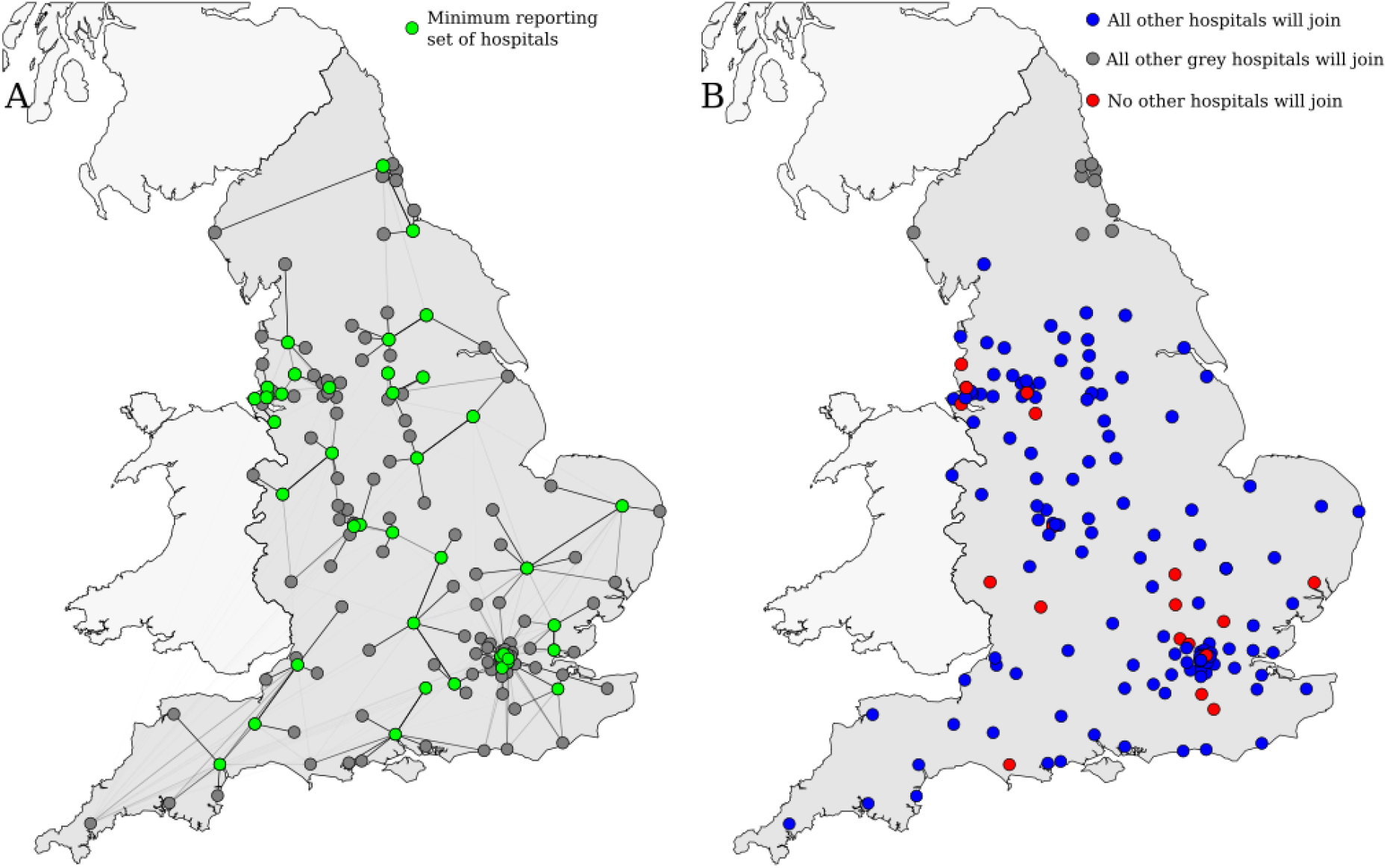
The geographical distribution of hospitals in the surveillance scheme. A) The minimal set of reporting hospitals needed to report on all hospitals, as found using the greedy algorithm. Green dots show the reporting set, grey dots the covered set and lines show the links over which patients previously discharged from other hospitals are included. B) The result of the snow-ball assumption (a hospital will start reporting once it is reported on) as a function of the first hospital to join the surveillance scheme. For the majority of hospitals (127/155), all other hospitals would join the scheme were they the first hospital to start reporting (blue dots). However, a small group in the North region (9/155) will only report on hospitals in the same region (grey dots), while for small number of hospitals (19/155) no others will join if they are the first in the surveillance system (red dots), because they do not receive over 1000 patients per year from any other single hospital, and hence no other hospitals will therefore be reported on and join the scheme.

## Discussion

To have the desired effect, incentives for hospitals to reduce their reported rates of AMR and other hospital-transmitted organisms need to align with the hospitals’ interests to reduce their numbers of colonised and infected patients. We show that this can be done by having hospitals report the number of cases among the patients they admit who have previously been discharged from other hospitals, as it separates the tasks of reporting and reducing incidence. In this way, hospitals report on the AMR incidence in other hospitals, not on their own incidence, and as a result they themselves do not suffer potential consequences from their reports. Additionally, if the recipient hospital is then rewarded for any case they find, a clear incentive is constructed to find as many cases as possible discharged from other hospitals, delivering a more reliable incidence estimate.

The proposed surveillance system intrinsically increases the number of covered hospitals. First and foremost, by reporting cases admitted after previously being discharged from other hospitals, not all hospitals need to participate for it to be possible to estimate incidence for all hospitals. In fact, a selected subset of only 26% of English hospitals resulted in enough patients admitted to another hospital within a year after discharge to estimate incidence in all hospitals in England. Even if hospitals join the surveillance system (the reporting set) at random, incidences for all hospitals can be obtained before all hospitals are reporting. The system therefore provides incidence estimates for more hospitals than participate. Furthermore, because the reported incidence for a certain hospital will often be the result of the pooled reports sent in by several other hospitals, the final measured incidence is less influenced by the screening rates of individual hospitals. The ranking of hospitals based on the agglomerated measurement can therefore be expected to be more robust than any measurement derived from single hospitals.

The number of hospitals participating in such a surveillance scheme could easily increase if hospitals were compensated for cases they find among patients admitted after having been discharged from another hospital, since there is no clear disadvantage to screening imported patients and reporting found cases. Subsequently, this effect may cause more hospitals to join: if a hospital’s incidence is reported by other hospitals, it may be inclined to start testing patients it admits after they have been discharged from other hospitals, if only to be able to compare incidences. Due to this snow-ball effect the system may not need to be mandatory, although a core group of participating hospitals may be desirable.

If the goal of reporting incidence changes from purely gathering information to creating incentives for improving performance by penalising hospitals with high incidences, either financially or reputationally, the proposed surveillance system still has value, because any repercussions associated with high incidence are incurred by a different hospital than the one that is screening patients. However, exactly which cases might be counted when penalising hospitals needs to be carefully considered. To promote information sharing between hospitals, only newly discovered AMR-positive patients should be used to determine penalties, and not those patients that were previously screened and labelled as carriers, to prevent the punishment of hospitals that actively try to share information about cases identified among their admitted patient population with other hospitals.

The proposed surveillance scheme exploits the structure of the hospital network, showing the added value of regarding hospitals as interconnected by shared patients instead of completely independent and isolated entities[8–12]. Previous studies have shown that patient sharing between hospitals significantly correlate with rates of Carbapenemase-Producing Enterobacteriaceae (CPE)[13], MRSA[14] or Clostridium difficile[15,16]. The influence of the hospital network formed by shared patients on the spread of hospital-associated pathogens has also been used to design early warning systems[16,17] or inform the distribution of resources for IPC[18], often reiterating the importance of centrally located hospitals. We present a novel viewpoint on using these hospital networks, by considering the interests of hospitals to report cases, thus actively using the shared patients to combat the spread of these pathogens.

### Limitations

The estimated incidence of a specific hospital measured by the reporting hospitals will not be identical to incidence measured within the specific hospital itself, because the readmitted patients are a specific subset of the original patient population and more likely carriers. However, readmitted populations will generally be broadly comparable between hospitals. Further, whilst this estimate may not precisely reflect the true incidence in a specific hospital, arguably neither does the self-reported rate. Comparing estimated incidences for hospitals with vastly different function, such as specialist hospitals, that have substantially different case-mix from the other hospitals, may need to be done carefully, for example using adjustment, as for standardised mortality rates.

We assumed that receiving hospitals are aware of patients’ previous hospital stays upon admission, to identify those that need to be screened. However, this may not necessarily be the case, in particular when the time since last discharge is relatively long. Reported incidences may therefore be slightly lower, because some shared patients might be missed. Although this would lower the surveillance system’s accuracy, the bias would be similar for all hospitals; in particular because multiple hospitals can report on each covered hospital, any inaccuracies on the single reporting hospital level will be averaged out.

We considered a cut-off for screening admissions of 1 year from previous discharge; in the general community, bacterial carriage may or may not persist over this period, making it harder to attribute colonisation status to the previous hospitalisation with confidence the longer a previous admission was in the past. This is particularly problematic if levels of community transmission start to exceed hospital-associated transmission. By shortening the cut-off time, the specificity of the surveillance system will increase, at the cost of its sensitivity. However, by recording all colonised patients who were previously admitted to another hospital, together with the time between admissions, it should be possible to estimate the relative contribution of community transmission to the importation of cases to all hospitals.

## Conclusion

We propose a new system to estimate incidences of AMR and other hospital-transmitted micro-organisms that does not rely on self-reporting, whereby instead surrounding hospitals report the incidence within the patient population admitted to their hospital who have recently being discharged from other hospitals. This decoupling of the hospital that is reporting from the hospital reported on is vital for delivering reliable incidence estimates, as it takes away the incentive to stop looking for cases by watching over the others. By reporting on other hospitals’ incidence, the surveillance scheme aligns financial and patient safety interests, encouraging hospitals to find and report as many cases as possible, making the surveillance scheme more resilient against ‘gaming’ and thus delivering a more robust comparison between hospitals.

## Supplementary information

Supplementary text: The game-theoretical implications of surveillance schemes

Supplementary table 1: Numbers of shared patients between hospitals, for cut-off time between admissions: one year, six months, three months, one month, and one week. Including list of hospital codes and names.

## Abbreviations

AMR: antimicrobial resistance
CPE: carbapenemase-producing enterobacteriaceae
HES: hospital episode statistics
IPC: infection prevention and control
MRSA: methicillin-resistant Staphylococcus aureus
NHS: National Health Service

## Authors’ contributions

TD, ASW, and JVR designed the study. TD and TS performed the analysis. KLH provided data. TD, TS, ASW, and JVR drafted the manuscript All authors revised the manuscript and gave final approval for publication.

## Availability of data and materials

The patient admission data were not collected for this study specifically, and the authors do not own the used datasets. Patient admission data from the English NHS HES (Hospital Episode Statistics. Copyright© 2015. Re-used with the permission of the Health and Social Care Information Centre. All rights reserved.) are available for researchers who meet the criteria for access to confidential data from NHS digital (digital.nhs.uk; Formerly Health and Social Care Information Centre). All other data needed to reproduce the analysis is provided in the supplementary information.

## References

1. Freeman T. Using performance indicators to improve health care quality in the public sector: a review of the literature. Heal Serv Manag Res. 2002;15:126–37.

2. Hibbard JH, Stockard J, Tusler M. Does publicizing hospital performance stimulate quality improvement efforts? Health Aff. 2003;22:84–94.

3. Maynard A, Bloor K. Will financial incentives and penalties improve hospital care? BMJ. 2010;340:c88.

4. Health Protection Agency. Quarterly Epidemiological Commentary: Mandatory MRSA, MSSA and E. coli bacteraemia, and C. difficile infection data (up to October – December 2012). 2013;1–8.

5. Johnson AP, Davies J, Guy R, Abernethy J, Sheridan E, Pearson A, et al. Mandatory surveillance of methicillin-resistant Staphylococcus aureus (MRSA) bacteraemia in England: the first 10 years. J Antimicrob Chemother. 2012;67:802–9.

6. Wertheim HF, Melles DC, Vos MC, van Leeuwen W, van Belkum A, Verbrugh H a, et al. The role of nasal carriage in Staphylococcus aureus infections. Lancet Infect Dis. 2005;5:751–62.

7. Calfee DP, Durbin LJ, Germanson TP, Toney DM, Smith EB, Farr BM. Spread of methicillin-resistant Staphylococcus aureus (MRSA) among household contacts of individuals with nosocomially acquired MRSA. Infect Control Hosp Epidemiol. 2003;24:422–6.

8. Donker T, Wallinga J, Grundmann H. Patient referral patterns and the spread of hospital-acquired infections through national health care networks. PLoS Comput Biol. 2010;6:e1000715.

9. Iwashyna TJ, Christie JD, Moody J, Kahn JM, Asch DA. The structure of critical care transfer networks. Med Care. NIH Public Access; 2009;47:787–93.

10. Nekkab N, Astagneau P, Temime L, Crépey P. Spread of hospital-acquired infections: A comparison of healthcare networks. PLOS Comput Biol. 2017;13:e1005666.

11. Huang SS, Avery TR, Song Y, Elkins KR, Nguyen CC, Nutter SK, et al. Quantifying interhospital patient sharing as a mechanism for infectious disease spread. Infect Control Hosp Epidemiol. 2010;31:1160–9.

12. Lee BY, McGlone SM, Song Y, Avery TR, Eubank S, Chang CC, et al. Social network analysis of patient sharing among hospitals in Orange County, California. Am J Public Health. 2011;101:707–13.

13. Ray MJ, Lin MY, Weinstein RA, Trick WE. Spread of Carbapenem-Resistant Enterobacteriaceae Among Illinois Healthcare Facilities: The Role of Patient Sharing. 2016;1–5.

14. Donker T, Wallinga J, Slack R, Grundmann H. Hospital networks and the dispersal of hospital-acquired pathogens by patient transfer. PLoS One. 2012;7:e35002.

15. Simmering JE, Polgreen L a., Campbell DR, Cavanaugh JE, Polgreen PM. Hospital Transfer Network Structure as a Risk Factor for Clostridium difficile Infection. Infect Control Hosp Epidemiol. 2015;1–7.

16. Fernández-Gracia J, Onnela J-P, Barnett ML, Eguíluz VM, Christakis NA. Influence of a patient transfer network of US inpatient facilities on the incidence of nosocomial infections. Sci Rep. 2017;7:2930.

17. Ciccolini M, Spoorenberg V, Geerlings SE, Prins JM, Grundmann H. Using an index-based approach to assess the population-level appropriateness of empirical antibiotic therapy. J Antimicrob Chemother. 2014;1–8.

18. Karkada UH, Adamic L a, Kahn JM, Iwashyna TJ. Limiting the spread of highly resistant hospital-acquired microorganisms via critical care transfers: a simulation study. Intensive Care Med. 2011;37:1633–40.

