## supplementary text for "Using hospital network-based surveillance for antimicrobial resistance as a more robust alternative to self-reporting"

*Supplementary material*

***The game-theoretical implications of surveillance schemes***

We can reduce the choices hospitals make to a game-theoretical framework, where each hospital has the choice to report or abstain from reporting; or equivalently test to try to identify carried resistance or not test. The pay-out for each choice is then based on the cost of the strategy (screening, isolation, etc.) and the resulting ranking (with high incidence hospitals fined and/or low incidence hospitals rewarded). For simplicity, we assume that abstaining from the reporting scheme results in a zero reported incidence.

**Self-reported surveillance scheme**

For a self-reported surveillance scheme, with two hospitals (A and B) with assumed equal AMR incidences, the cost build up would be as follows:

- Cost of screening and isolation: C

- Reward for lowest incidence (Winning the ranking): W

- Fine for highest incidence (Losing the ranking): L

The pay-out matrix for two hospitals can then be constructed using any cost as negative pay-out and rewards or compensations as positive pay-out. They then look as follows, with each cell giving the pay-outs for {A,B}:

|  |  | B | |
| --- | --- | --- | --- |
|  | A\**B** | Abstain | Report |
| A | Abstain | 0,**0** | W**, - C - L** |
|  | Report | - C- L, **W** | - C, -**C** |

As long as all of the cost parameters are positive, both hospitals will always choose to abstain from reporting, as - C - L < 0 and - C < W.

If we assume that one of the hospitals has a higher incidence (hospital B), the pay-out matrix becomes asymmetrical. With hospital B losing the ranking more often and the cost of screening being different for the two hospitals (C_A_ and C_B_) the matrix becomes symmetric:

|  |  | B | |
| --- | --- | --- | --- |
|  | A\**B** | Abstain | Report |
| A | Abstain | 0,**0** | W, **- C_B_ - L** |
|  | Report | **-** C_A_ - L, **W** | W **-** C_A_, **- C_B_ - L** |

Although the exact values change, this does not change the stable strategy (abstain/abstain).

**Proposed reporting scheme**

Using the proposed surveillance scheme, the result of the ranking changes place, with reporting being associated with winning the ranking.

|  |  | B | |
| --- | --- | --- | --- |
|  | A\**B** | Abstain | Report |
| A | Abstain | 0,**0** | - L, **W – C** |
|  | Report | W - C, **- L** | - C, **- C** |

Both hospitals will now keep reporting if the cost of screening and isolating is lower than the fine for losing the ranking (C < L). Moreover, if the reward for winning the ranking is higher than the cost of screening and isolation (W > C), starting screening is the best option, making report/report the only stable equilibrium.

|  |  | B | |
| --- | --- | --- | --- |
|  | A\**B** | Abstain | Report |
| A | Abstain | 0,**0** | - L, **W - C_B_** |
|  | Report | W - C_A_, **L** | W - C_A_, **- C_B_ - L** |

If hospital B has a higher incidence it will stop screening if hospital A is screening, because it does not gain an advantage in the ranking by screening imported cases from the other hospital (but can reduce its own total cost by not screening).

|  |  | B | |
| --- | --- | --- | --- |
|  | A\**B** | Abstain | Report |
| A | Abstain | 0,**0** | - L, **+ W** |
|  | Report | + W, **- L** | + W, **- L** |

If hospitals are compensated for the cost of screening and isolation, hospital B will be ambivalent about starting or stopping screening if hospital A is screening, but will start screening if hospital A is not.

|  |  | B | |
| --- | --- | --- | --- |
|  | A\**B** | Abstain | Report |
| A | Abstain | 0,**0** | - L, **W + C_B_** |
|  | Report | W + C_A_, **- L** | W + C_A_, **C_B_ - L** |

However, if hospitals get a reward per case that is higher than its cost of isolation (denoted as adding instead of subtracting C_A/B_), hospital B will keep screening because C_B_-L >-L.

Under first scenario, without a priori information about incidence in each hospital, the problem can be view as a classical prisoner’s dilemma: the least costly strategy for all hospitals is to abstain from reporting, whereas the individual hospitals’ best strategy is to report. However, if differences in incidence are taken into account, it may not always be beneficial for the hospital with highest incidence to start screening, as it cannot win the ranking. It should however be noted that in a full hospital network with more than two hospitals, this strategy would only apply to the hospital with the highest incidence (if it was fully aware of its all hospitals’ positions in the ranking). The second worse hospital is still theoretically able to improve its relative ranking by reporting. In practice, the drive for hospitals to participate in the proposed surveillance scheme will probably be a function of the difference between the true and reported ranking and the rewards and/or fines associated with the ranking.
